# *HybridExpress*: an R/Bioconductor package for comparative transcriptomic analyses of hybrids and their progenitors

**DOI:** 10.1101/2024.04.02.587701

**Authors:** Fabricio Almeida-Silva, Lucas Prost-Boxoen, Yves Van de Peer

## Abstract

Hybridization, the process of crossing individuals from diverse genetic backgrounds, plays a pivotal role in evolution, biological invasiveness, and crop breeding. At the transcriptional level, hybridization often leads to complex non-additive effects, presenting challenges for understanding its consequences. Although standard transcriptomic analyses exist to compare hybrids to their progenitors, such analyses have not been implemented in a software package, hindering reproducibility. Here, we introduce *HybridExpress*, an R/Bioconductor package designed to facilitate the analysis, visualization, and comparison of gene expression patterns in hybrid triplets (hybrids and their progenitors). *HybridExpress* provides users with a user-friendly and comprehensive workflow that includes all standard comparative analyses steps, including data normalization, calculation of midparent expression values, sample clustering, expression-based gene classification into categories and classes, and overrepresentation analysis for functional terms. We illustrate the utility of *HybridExpress* through comparative transcriptomic analyses of cotton allopolyploidization and rice root trait heterosis. *HybridExpress* is designed to streamline comparative transcriptomic studies of hybrid triplets, advancing our understanding of evolutionary dynamics in allopolyploids, and enhancing plant breeding strategies. *HybridExpress* is freely accessible from Bioconductor (https://bioconductor.org/packages/HybridExpress) and its source code is available on GitHub (https://github.com/almeidasilvaf/HybridExpress).

## Introduction

Hybrids result from crossing individuals from different species, subspecies, or genetic lineages. Hybridization plays a crucial role in evolution (Abbott *et al*., 2013; Taylor & Larson, 2019), biological invasiveness (Ellstrand & Schierenbeck, 2000), and crop breeding through heterosis (Fu *et al*., 2014; Labroo *et al*., 2021). Hybrids, whether homoploid or allopolyploid, combine two distinct genomes and their associated regulatory networks in a single nucleus, resulting in potential incompatibilities (Landry *et al*., 2007; Maheshwari & Barbash, 2011; Sloan *et al*., 2023), (epi)genomic changes (Baack & Rieseberg, 2007; Greaves *et al*., 2015), and phenotypic novelties (Dittrich-Reed & Fitzpatrick, 2013). Hybridization can also lead to complex and unpredictable alterations of transcriptional regulation, a phenomenon known as “transcriptomic shock” (Hegarty *et al*., 2006; Buggs *et al*., 2011).

Comparative transcriptomics studies of hybrid triplets (i.e., hybrids and their parent species) have provided novel insights on changes in transcriptional regulation following genome merging. These transcriptomic responses typically manifest as non-additive expression patterns in hybrids, such as expression-level dominance (i.e., expression in the hybrid mimics that of one of its progenitors) and transgressive up- and down-regulation (i.e., expression is higher or lower in the hybrid relative to both progenitors) (Osborn *et al*., 2003; Adams *et al*., 2003; Otto, 2003; Adams & Wendel, 2005; Chen, 2007; Yoo *et al*., 2013, 2014). Such changes have been documented in several plant species, including maize (Guo *et al*., 2004, 2006; Auger *et al*., 2005; Stupar *et al*., 2008), rice (Wei *et al*., 2009; He *et al*., 2010), cotton (Rapp *et al*., 2009; Flagel & Wendel, 2010), *Arabidopsis* sp. (Wang *et al*., 2006), Monkeyflower (Edger *et al*., 2017), and *Senecio sp*. (Hegarty *et al*., 2008), as well as in animals, such as insects (Wang *et al*., 2015) and fish (Gao *et al*., 2013; Li *et al*., 2018). Many factors can influence the transcriptomic changes in hybrids, including species-specific characteristics, the nature of the cross, environmental conditions (Bardil *et al*., 2011; Shimizu-Inatsugi *et al*., 2017), and tissue specificity (Li *et al*., 2014, 2020).

To increase our understanding of the transcriptome-level effects of hybridization, methods for comparing, analyzing, and visualizing gene expression patterns in hybrids have been developed and standardized. These methods include normalization techniques to account for differences in biomass (for polyploids) and library size (Visger *et al*., 2019; Coate, 2023), and the classification of genes in classes and categories based on their expression patterns (Hegarty *et al*., 2006; Flagel *et al*., 2008; Hovav *et al*., 2008; Rapp *et al*., 2009). Notably, Rapp et al. (2009) were, to our knowledge, pioneers in introducing a twelve-category classification for gene expression, which has since become a standard approach in comparative transcriptomics of hybrid triplets (see, for examples, Bardil et al., 2011; Chagué et al., 2010; Chelaifa et al., 2010; Wu et al., 2018; Yoo et al., 2013). However, these methods have not been implemented in a user-friendly software package, which poses a challenge for novice bioinformaticians, and hinders reproducibility and efficiency.

Here, we present *HybridExpress*, an R/Bioconductor package that provides a comprehensive and user-friendly framework to analyze, visualize, and compare gene expression patterns in hybrid triplets. From gene expression data, *HybridExpress* allows users to normalize data, calculate midparent expression values, explore sample clustering, classify genes into expression-based categories and classes (as per Rapp et al., 2009), and perform overrepresentation analysis for functional terms. By applying *HybridExpress* to real data sets, we demonstrate how the package can be used to better understand the transcriptomic effects of allopolyploidization in cotton and root trait heterosis in rice. *HybridExpress* will likely be a central resource in comparative transcriptomic analyses of hybrid triplets, from evolutionary applications to plant breeding.

## Materials and Methods

### Implementation

For seamless integration with other Bioconductor packages, input data for *HybridExpress* are either base R or core Bioconductor S4 classes (e.g., *SummarizedExperiment* objects for quantitative data and associated metadata) (Morgan *et al*., 2023). For increased efficiency, functions that perform operations on large sets (e.g., overrepresentation analyses) can be parallelized using the back-ends provided by the BiocParallel package (Morgan *et al*., 2014).

### Calculating (in silico) midparent expression values

Midparent expression values are typically used as a reference point to test for genetic additivity, and deviations from midparent values are interpreted as resulting from non-additive effects (Gianinetti, 2013). By default, *HybridExpress* creates *in silico* samples with midparent values by calculating the mean expression of sample pairs containing a sample from each parent. Users can also calculate the weighted mean of sample pairs, which can help account for differences in ploidy levels in newly synthesized polyploids (Gianinetti, 2013).

### Data normalization

By default, *HybridExpress* normalizes count data by library size using size factors estimated with the ‘median of ratios’ method implemented in the DESeq2 package (Anders & Huber, 2010; Love *et al*., 2014). This technique corrects for differences in sequencing depth across samples, and it assumes equal amounts of mRNA per cell between conditions (Evans *et al*., 2018). However, when this assumption is violated (*e*.*g*., when comparing individuals with different ploidy levels and, hence, different cell sizes and/or biomass/genome ratio), users can also normalize data with spike-in controls that aim at correcting for differences in cell size and units of biomass per genome (Evans *et al*., 2018; Coate, 2023).

### Exploratory data analysis

To help users detect potential issues in sample grouping (*e*.*g*., batch effects and high heterogeneity), graphical functions are available to easily create heatmaps of hierarchically clustered samples (from pairwise correlations), and principal component analysis plots. For both analyses, expression data are filtered to include only the top *N* genes with the highest variances to reduce noise. Then, a variance-stabilizing transformation is applied to remove the mean-variance relationship typically observed in RNA-seq count data. All figures generated with *HybridExpress* are publication-ready, and they can be further customized with the ggplot2/grid plotting system (Wickham, 2011).

### Identification of differentially expressed genes (DEGs)

Differentially expressed genes are detected using the DESeq2 algorithm (Love *et al*., 2014) on four contrasts: parent 2 *versus* parent 1; hybrid *versus* parent 1; hybrid *versus* parent 2; and hybrid *versus* midparent value. The function *get_deg_list()* returns gene-wise test statistics for each contrast, and the function *get_deg_counts()* summarizes results in a table with frequencies (absolute and relative) of up- and down-regulated genes per contrast. Frequencies of DEGs in all contrasts can be summarized in a single figure with the function *plot_expression_triangle()*, which creates a DEG triangle commonly used in publications (Rapp *et al*., 2009).

### Expression-based gene classification

From the gene-wise test statistics returned by *get_deg_list()*, the function *expression_partitioning()* can be used to classify genes into one of the twelve expression-based categories proposed by Rapp et al. (2009). Besides, since many of such categories display biologically redundant information, genes can be further grouped into five major classes, namely: i. transgressive up-regulation; ii. transgressive down-regulation; iii. additivity; iv. expression-level dominance towards parent 1; and expression-level dominance towards parent 2. Results can be graphically summarized in publication-ready figures with the functions *plot_expression_partitions()* and *plot_partition_frequencies()*.

### Functional analyses

Overrepresented functional terms (e.g., biological processes or metabolic pathways) among expression-based classes and categories can be identified with the *ora()* function. Functional annotation for the *ora()* function can be passed as a two-column data frame with genes and the functional terms associated with them, giving users the flexibility to use functional annotation from any database, as well as custom annotation (e.g., experimentally validated genes).

### Benchmark data sets

Data for benchmarks 1 and 2 were obtained from Dong *et al*. (2022) and Zhai *et al*. (2013), respectively. In the first benchmark data set, authors investigated the transcriptomic responses to modest salinity stress in two allotetraploid cotton species (*Gossypium hirsutum* and *G. mustelinum*, AD genome) relative to their model diploid progenitors (A and D genomes). For each species, three biological replicates of 2-week-old seedlings were obtained in control and stress conditions. In the second benchmark data set, authors sequenced the root transcriptomes of the super-hybrid rice (*Oryza sativa*) variety Xieyou 9308 and its parents (lines R9308 and Xieqingzao B) during heading and tillering stages, with two replicates for each line and stage. The goal of the study was to understand the transcriptomic basis of heterosis (i.e., superior performance of hybrids compared to parents) in root traits. Functional annotation (Gene Ontology, InterPro domains, and MapMan pathways) for the cotton and rice genomes were obtained from PLAZA Dicots 5.0 and PLAZA Monocots 5.0, respectively (Van Bel *et al*., 2022).

## Results and Discussion

### Use case 1: transcriptional responses to salt stress in allopolyploid cotton species

We used *HybridExpress* to perform comparative transcriptomic analyses in two allotetraploid cotton species (*Gossypium hirsutum* and *G. mustelinum*, hereafter referred to as ‘AD1’ and ‘AD4’, respectively) relative to their diploid progenitors (A2 and D5) under moderate salinity stress (see Materials and Methods for details). We processed the original count matrix to remove lowly expressed genes (sum of counts<10), include midparent expression values, and normalize data by library size. Principal component analyses revealed that samples grouped well by species and treatment, but an outlier was detected and removed from further analyses (Fig. 1A and 1B). Importantly, such outlier was also detected in the original publication, highlighting that results are consistent (Dong *et al*., 2022). Principal component 1, which describes most of the variation (42.3% and 47.2% for AD1 and AD4, respectively), separates samples by species, and it shows that both allopolyploids are closer to the A2 parent than to the D5 parent (Fig. 1A and 1B).

**Fig. 1.**
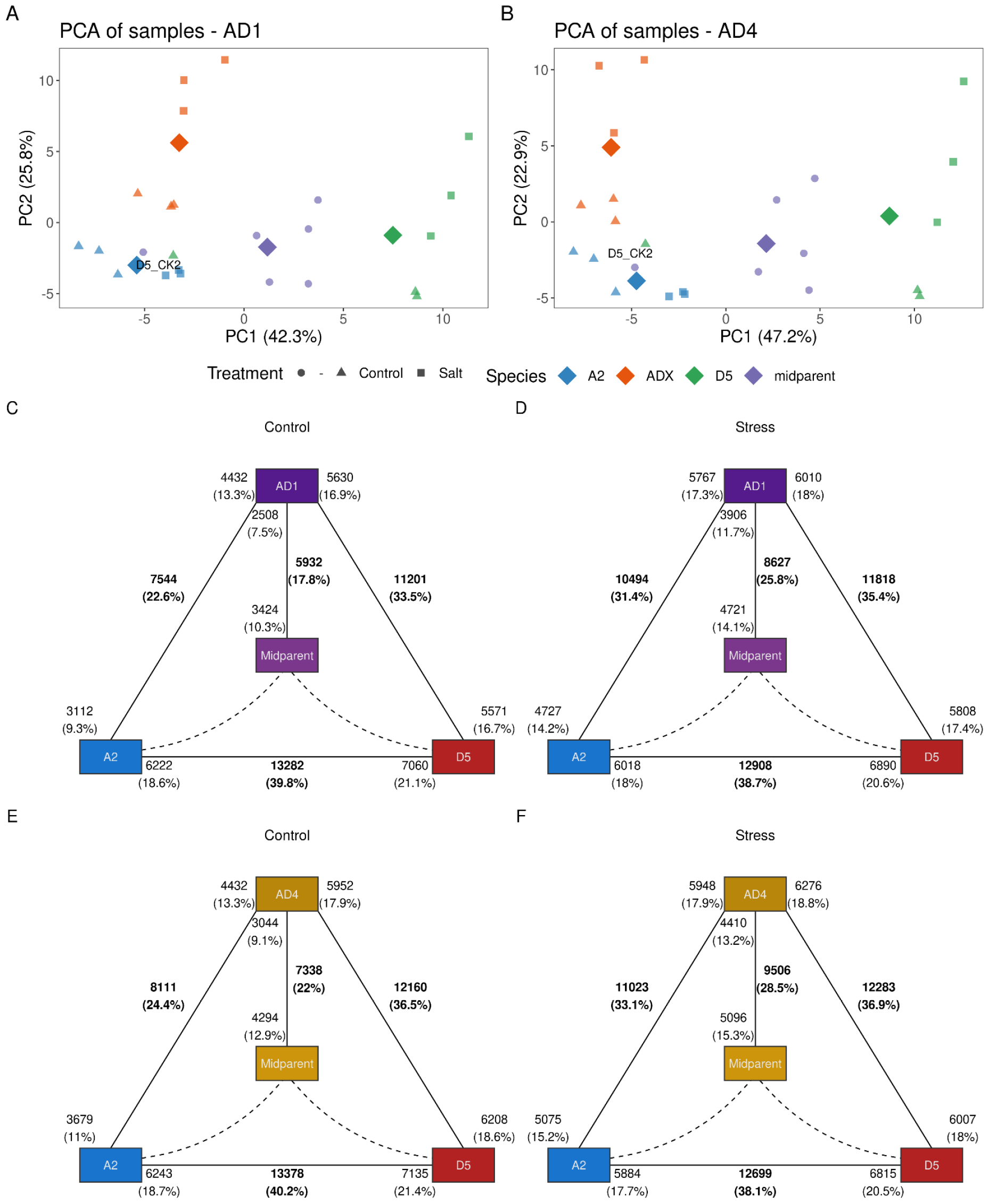
Sample grouping and frequencies of differentially expressed genes (DEGs) between species pairs for allopolyploid cotton (see text for details). **A-B**. Principal component analysis of samples for the hybrid triplet including genomes AD1 (*Gossypium hirsutum*) and AD4 (*Gossypium mustelinum*). Figures were created with the *pca_plot()* function. Samples group well by species and conditions, but one outlier (sample *D5_CK2*) can be detected. Both allotetraploids are closer to the A2 parent than to the D5 parent. **C-F**. Expression triangle plot showing the frequencies of DEGs for each pairwise species pair, with midparent value (*in silico*) in the center, for AD1 (control and stress) and AD4 (control and stress). Figures were created with the *plot_expression_triangle()* function. Figures show that the number of DEGs in allotetraploids relative to the D5 parent is greater than the number of DEGs relative to the A2 parent, indicating expression-level dominance towards A2. However, the dominance is greatly reduced under salt stress.

We further identified differentially expressed genes (DEGs) between species pairs for each triplet and condition. For both allotetraploids, the number of DEGs relative to the D5 parent is greater than the number of DEGs relative to the A2 parent, indicating an expression-level dominance (ELD) towards A2. However, the difference in number of DEGs is reduced upon salt stress (Fig. 1C to 1F), suggesting that the stress condition disrupts ELD. When exploring the overlap of DEGs in allotetraploids relative to both parents in control and stress conditions, we found many shared DEGs relative to both parents and conditions (*N* = 2187 and 2693 for AD1 and AD4, respectively) (Fig. 2A and 2B). Besides, most DEGs relative to the D5 parent are shared between control and stress conditions, while DEGs relative to the A2 parent are mostly specific to each condition (Fig. 2A and 2B).

**Fig. 2.**
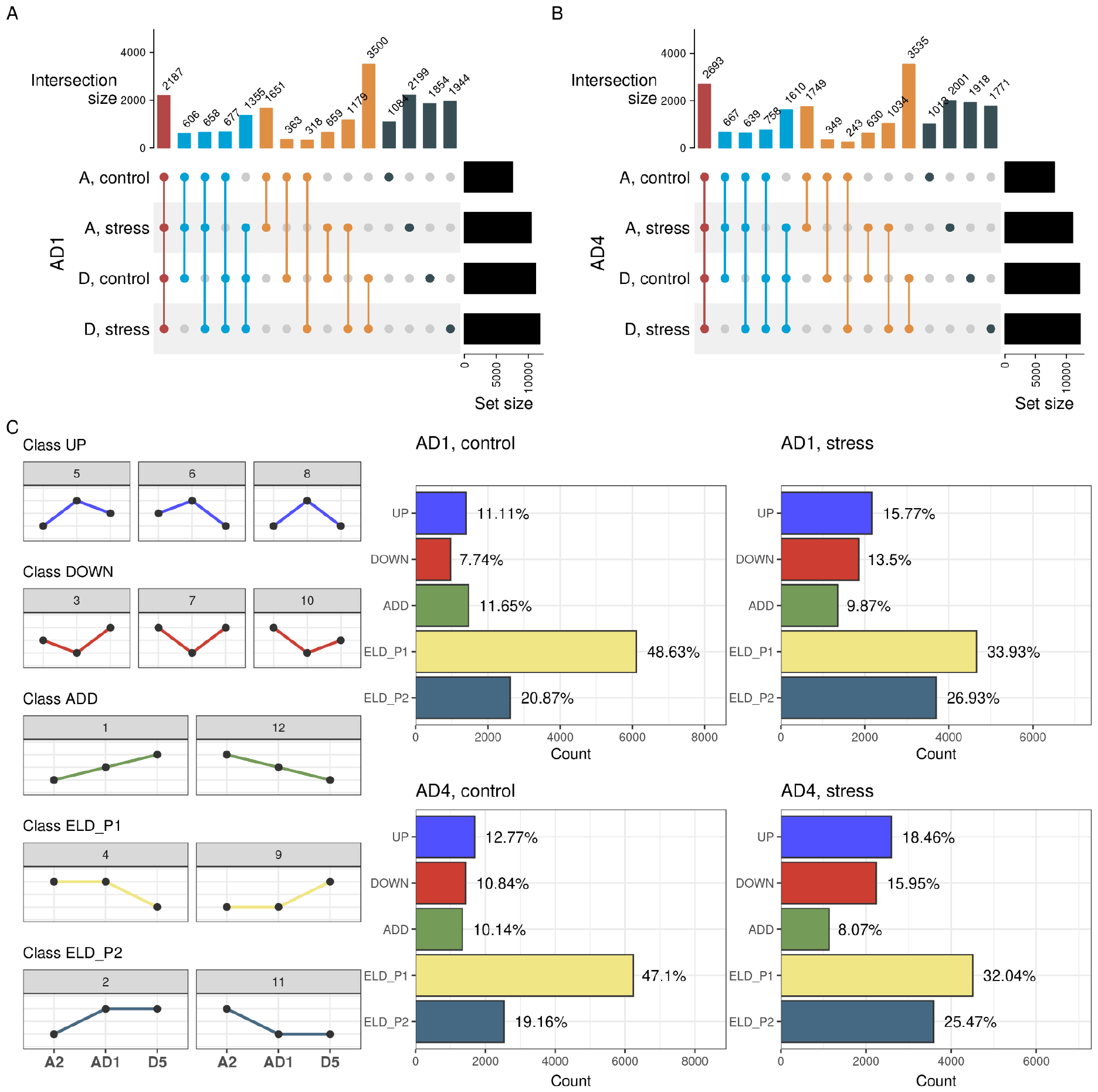
Patterns of shared DEGs in cotton allotetraploids, and expression-based partitioning of genes in classes. **A-B**. UpSet plots showing the overlaps of DEGs in the allotetraploids relative to both parents in control and stress conditions. Different colors indicate different overlap sizes (i.e., shared by all four sets, three sets, two sets, and set-specific DEGs). **C**. Frequency of genes per expression-based class for AD1 (control and stress) and AD4 (control and stress). Line plots on the left are schemes to visually exemplify what the expression patterns of genes in each class and category look like. Most genes display expression-level dominance towards parent 1 (A2), but the dominance is greatly reduced upon stress.

To better understand global expression patterns in allotetraploids relative to their parents, we classified genes into expression-based classes and calculated the frequency of genes in each class. For both tetraploids, we observed that most genes display ELD towards the A2 parent in control and stress conditions, but such bias towards the A2 parent is greatly reduced under salt stress (Fig. 2C). This finding is in line with what we observed based on the number of DEGs alone (Fig. 1C to 1F), providing strong evidence for a salt stress-mediated disruption of ELD. Genes displaying ELD towards the A2 parent were associated with photosynthesis, ribosome biogenesis, RNA modifications, and circadian rhythm, while those displaying ELD towards the D5 parent were associated with phenylpropanoid biosynthesis, and regulation of defense response (e.g., ERF and WRKY transcription factors) (Supplementary Table S1). Under salt stress, genes displaying transgressive up-regulation were involved in abscisic acid binding, response to salicylic acid, and redox metabolism. Genes displaying transgressive down-regulation under salt stress were involved in cell wall organization, such as cellulose synthases, galacturonan metabolism, and endoglucanases.

Considering the spatiotemporal dynamics of transcriptional regulation, the effects of allopolyploidization at the transcriptional level are likely to vary in different conditions, tissues, and species. While Flagel & Wendel (2010) observed biased expression towards the D genome in petal transcriptomes of five natural *Gossypium* allotetraploids, we observed biased expression towards the A genome in seedlings, suggesting that tissue-specific transcriptional programs play a role in expression-level dominance. Besides, while we observed reduced ELD under salt stress, Dong *et al*., (2022) found that salt stress increased homoeolog expression bias (i.e., preferential expression of one homoeolog relative to the other), highlighting that expression-level dominance (or genome expression dominance) and homoeolog expression bias are different phenomena (Grover *et al*., 2012). Collectively, our findings indicate how *HybridExpress* can help understand the transcriptomic effect of allopolyploidization.

### Use case 2: heterosis in rice root traits

To understand the transcriptomic basis of heterosis in root traits in rice (*Oryza sativa* ssp. *japonica*), we compared transcriptomes of the super-hybrid rice variety Xieyou 9308 (hereafter referred to as ‘F1’) with its parents (lines R9308 and Xieqingzao B, hereafter referred to as ‘P1’ and ‘P2’) at heading and tillering stages (see Materials and Methods for details). As we did for the cotton data set, we filtered out lowly expressed genes (sum of counts <10), calculated midparent expression values, and normalized data by library size. Principal component analyses revealed that samples from the same line clustered together as expected, but a large within-group variance was observed for hybrid samples at tillering stage (Fig. 3A and 3B). Interestingly, the hybrid line was closer to P1 at the heading stage, but closer to P2 at the tillering stage, suggesting a shift in ELD at different developmental stages (Fig. 3A and 3B).

**Fig. 3.**
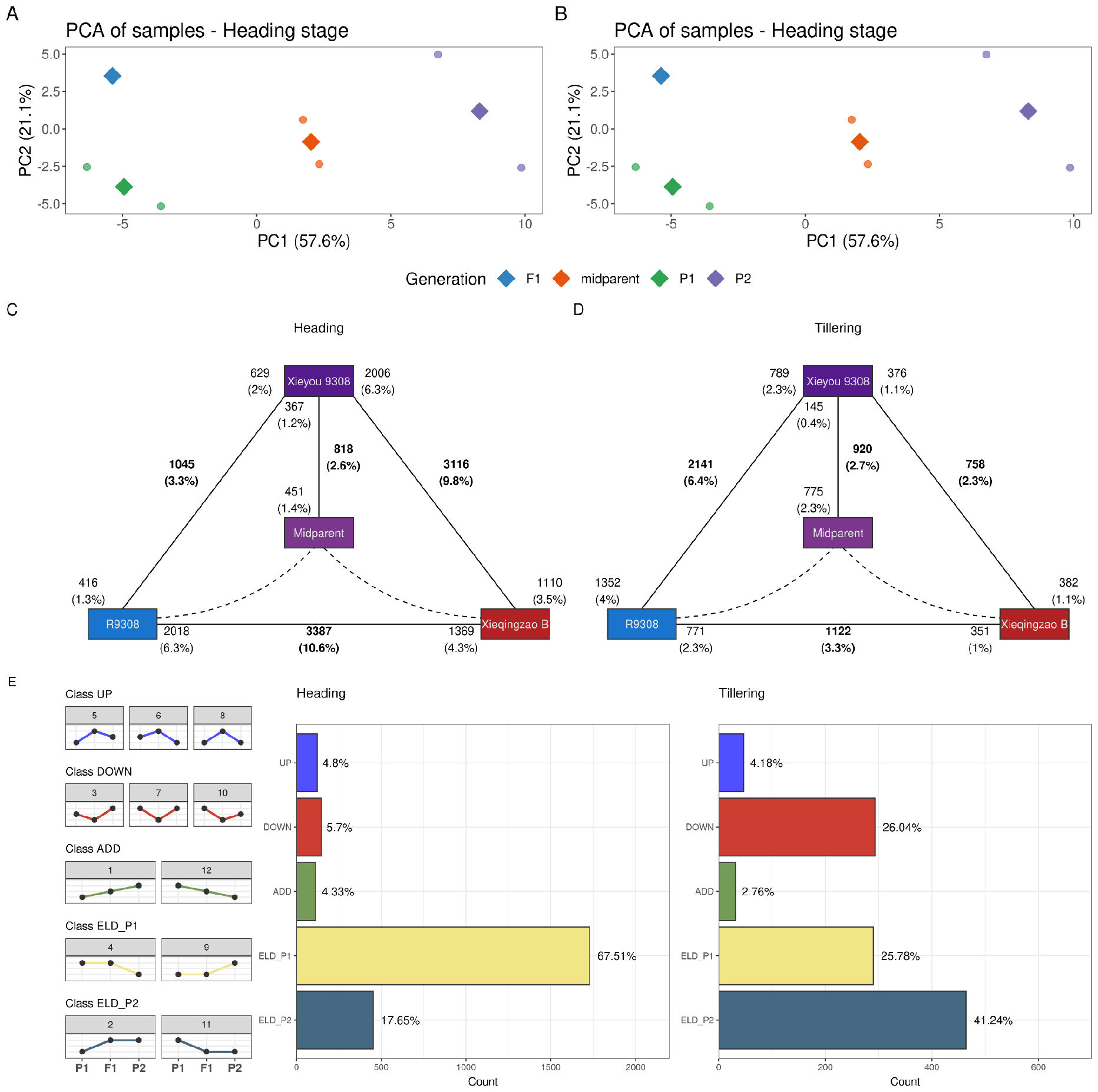
Comparative analyses of rice root transcriptomes in a hybrid triplet displaying heterosis of root traits. **A-B**. Principal component analysis of samples at heading and tillering stages. Figures were created with the *pca_plot()* function. The hybrid line is closer to P1 (line R9308) at the heading stage, but closer to P2 (line Xieqingzao B) at the tillering stage. **C-D**. Expression triangle plots showing the frequencies of DEGs for each pairwise combination of lines, with midparent values (*in silico*) in the center, at heading and tillering stages. Figures were created with the *plot_expression_triangle()* function. Figures show that the number of DEGs in the hybrid relative to P2 is greater at the heading stage, while the opposite trend is observed at the tillering stage. **E**. Frequency of genes per expression-based class at heading and tillering stages. Most genes display expression-level dominance towards P1 at the heading stage, while most genes display expression-level dominance towards P2 at the tillering stage, indicating a shift in dominance between developmental conditions.

Next, we identified DEGs between pairwise combinations of lines during both developmental stages. We found a greater number of DEGs relative to P2 at the heading stage, but the opposite trend at the tillering stage, again suggesting a developmental stage-mediated shift in ELD (Fig. 3C and 3D). To validate such a shift, we classified genes in expression-based classes and calculated the frequency of genes per class in both developmental stages. Most of the genes (67.51%) displayed ELD towards P1 at the heading stage, while most of the genes (41.24%) displayed ELD towards P2 at the tillering stage (Fig. 3E). However, a large but smaller fraction of the genes also displayed transgressive down-regulation and ELD towards P1 at the tillering stage (Fig. 3E). Genes displaying transgressive down-regulation were associated with flavone biosynthesis, redox metabolism, MYB transcription factors, and CASP-like proteins 1U (Supplementary Table S2). Genes displaying ELD towards P1 at heading stage were enriched in aquaporins and diacylglycerol kinases, while genes displaying ELD towards P2 at tillering stage were associated with apoptosis-activating factors, gibberellin biosynthesis, and terpene synthases. Collectively, our findings provide strong support for a developmental stage-mediated shift in genomic expression dominance in this rice hybrid triplet.

Changes in gene expression have been linked to heterosis in many plant traits, such as seed germination and plant height in maize (Birdseye *et al*., 2021; Wan *et al*., 2022), yield in rice (Sun *et al*., 2023), and nicotine biosynthesis in *Nicotiana tabacum* (Tian *et al*., 2018). However, such changes in transcriptional programs can (and possibly will) vary in a spatiotemporal scale, resulting in tissue- or condition-specific changes in the strength of heterosis (Ko *et al*., 2016; Yang *et al*., 2021). For instance, Zhai *et al*. (2013) demonstrated that heterosis in root traits (e.g., root-to-shoot ratio and root dry weight) is stronger during the tillering stage, which is in line with the more dramatic non-additive transcriptomic effects we observed in this stage. In particular, the large frequencies of genes displaying expression-level dominance and transgressive down-regulation at the tillering stage compared to the heading stage could explain the stronger heterotic effects at the phenotypic level, highlighting the potential of *HybridExpress* in understanding the molecular mechanisms underlying trait heterosis.

## Conclusion

In this study, we described *HybridExpress*, an R/Bioconductor package that provides researchers with a comprehensive and easy-to-use framework for comparative transcriptomic analyses of hybrids relative to their progenitors. Features included in *HybridExpress* allow users to perform a complete comparative transcriptomics workflow, including data processing, identification of differentially expressed genes, expression-based classification of genes, and functional analyses. Benchmarks demonstrated how *HybridExpress* can be used to better understand the transcriptomic effects of allopolyploidization and heterosis.

## Acknowledgements

YVdP acknowledges funding from the European Research Council (ERC) under the European Union’s Horizon 2020 research and innovation program (No. 833522). YVdP and FA-S acknowledge funding from Ghent University (Methusalem funding, BOF.MET.2021.0005.01). LP-B acknowledges Fonds Wetenschappelijk Onderzoek (FWO, 11H0426N) of Flanders for a PhD scholarship.

## Author contributions

FA-S, LP-B, and YVdP conceived the ideas and designed the methodology. FA-S and LP-B collected the data. FA-S analyzed the data. FA-S, LP-B, and YVdP led the writing of the manuscript. All authors contributed critically to the drafts and gave final approval for publication.

## Data availability

To ensure full reproducibility, all code and data used in this manuscript are available in a GitHub repository at https://github.com/almeidasilvaf/HybridExpress_paper, and an online Quarto book with executed code chunks and output are available at https://almeidasilvaf.github.io/HybridExpress_paper/.

